# Statistical Mechanics of Multistable Perception

**DOI:** 10.1101/008177

**Authors:** Gurinder S. Atwal

**Affiliations:** Cold Spring Harbor Laboratory, Cold Spring Harbor, NY 11724

## Abstract

The stochastic dynamics of multistable perception poses an enduring challenge to our understanding of neural signal processing in the brain. We show that the emergence of perception switching and stability can be understood using principles of probabilistic Bayesian inference where the prior temporal expectations are matched to a scale-free power spectrum, characteristic of fluctuations in the natural environment. The optimal percept dynamics are inferred by an exact mapping of the statistical estimation problem to the motion of a dissipative quantum particle in a multi-well potential. In the bistable case the problem is further mapped to a long-ranged Ising model. Optimal inference in the presence of a 1/f noise prior leads to critical dynamics, exhibiting a dynamical phase transition from unstable perception to stable perception, as demonstrated in recent experiments. The effect of stimulus fluctuations and perception bias is also discussed.

---

Multistable perception is a striking quantitative feature of human cognition and has stimulated much research in neuroscience and psychophysics [1, 2]. A classic example is the Necker cube whereby our visual perception of the ambiguous figure switches stochastically between two interpretations even though the image is static [3]. Stochastic percept switching in vision, and other sensory modalities [4], provides a useful experimental tool for investigating perception organization and neural information processing. Recent investigations have also demonstrated that, under differing experimental conditions such as intermittent stimulus removal, the switching rate can slow down and can even come to a stop, indicating a dynamical phase transition from unstable perception to stable perception [5, 6].

The mechanistic neural basis underlying multistable perception is not well understood. Most computational models assume that the biological origin of switching is either due to inherent neural noise, adaptation processes or optimal signal processing (see, for example [7–10]). However, these models are unable to easily account for the reported transitions to perception stabilization and it is unclear what role evolution in the natural environment has played. Here it is shown that, by framing perception as a unified Bayesian inference task, both perception switching and stability emerge naturally from rational and optimal interpretations computed by the brain, without the need to invoke *ad hoc* neurophysiological processes such as neural noise. The alternate percepts represent hypotheses of the external world, determined by a tradeoff between the sensory data and *a priori* beliefs. Central to our formalism is that the prior expectations are matched to the observed scale-free statistics of natural temporal patterns. Remarkably, the Bayesian formulation of the inference problem can be mapped to the dynamics of a dissipative quantum system, allowing us to deploy the powerful tools of statistical physics to understand the emergence of dynamical phase transitions in statistical learning of complex data.

We start by assuming that the input data comprises of *N* measurements *x*_1_*x*_2_…*x*_*N*_ ≡ {*x*_*t*_} at discrete times *t* over a duration time *T*. The data arises probabilistically from a model parameterized by *α*, representing the feature or percept, and the perception task is to estimate the time sequence {*α*_*t*_}. The input sequence is taken to be conditionally independent i.e.

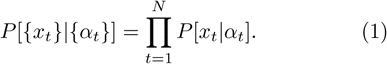

If {*x*_*t*_} arises unambiguously from a single true model 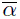 then it is expected that our estimate *α*(*t*) will fluctuate close to 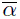 [11]. It is envisioned that there is an energy landscape in the *α*-space with a minimum at 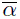 and the optimal estimate of *α*(*t*) will tend to keep it close to the ground state.

A more interesting, and generalized, question arises when we consider a non-trivial form of the extended energy landscape. In particular, there may be *C* ≥ 2 plausible interpretations consistent with the input data. It is useful to rewrite Eq. (1) using the following identity

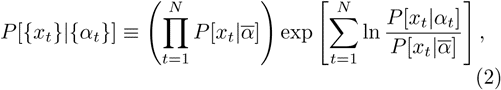

where 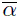 corresponds to any one of the *C* stable percepts. If the time variation of *α*(*t*) is slow, we effectively collect many samples of *x* before *α* changes, allowing us to replace the sum over samples by its continuum limit:

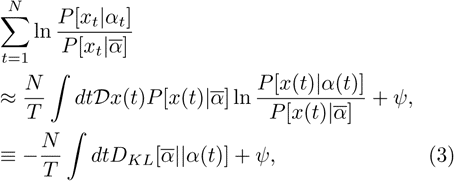

where *D*_*K L*_ is the Kullback-Leibler divergence between two distributions [12], and *ψ* are the sampling fluctuations in the data away from the thermodynamic limit, obeying the asymptotic constraint *ψ* → 0 as *N* → ∞. The *D*_*K L*_ term plays the role of a potential energy *V* [*α*(*t*)] with minima located at the set 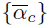.

Bayes’ theorem then furnishes us with the posterior probability distribution of the sequence of model parameters given the input data,

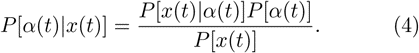

In multistable perception phenomena the incoming data *x*(*t*) is ideally constant, but in practise there may be fluctuations in the input which we will discuss later. To proceed further we require *a priori* hypotheses about how *α*(*t*) can vary in time. To represent our prior expectation that the local dynamics of *α*(*t*) vary slowly, we constrain the time-averaged square of the time-derivative, penalizing rapid changes in the estimate of *α*(*t*) [10],

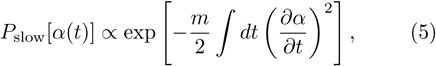

where *m* is a constant, weighting the strength of this prior. This prior serves to regularize the statistical model and prevents overfitting to the input data. Another, more general, prior belief is that the dynamics of the external world are drawn from a distribution with a scale-invariant power spectrum, as ubiquitously observed from a wide range of physical and biological settings [13–16]. This encapsulates the idea that biological systems have evolved to respond optimally to physical properties of natural environments. In frequency coordinates, the simplest form of the general environmental prior is

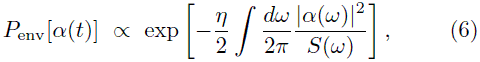

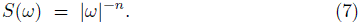

The parameter *η* determines the strength of prior belief and *n* is the scaling dimension of the power spectrum. When 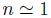 we have the familiar “1/f noise”, also known as flicker noise or pink noise. We stress that the noise here refers to the statistics of the environment and not the brain, which we assume to be noiseless. Taken together, the priors Eq. (5) and Eq. (6) describe fluctuations in *α*(*t*) which are *ω*^−2+(2−*n*)*θ*(2−*n*)^ up to a cross-over frequency, *ω*_*c*_ ∼ (*η*/*m*)^2−*n*^. Combining both priors in the Bayesian posterior probability Eq. (4) we obtain a unified path-integral expression

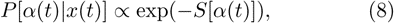

where the action *S*[*α*(*t*)] is given by

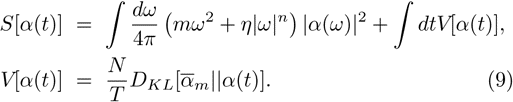

In the time-domain,

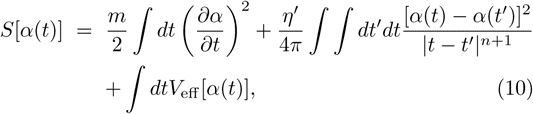

where we have made use of the identity 2*α*(*t*)*α*(*t*′) ≡ [*α*^2^(*t*) + *α*^2^(*t*′)] − [*α*(*t*) − *α*(*t*′)]^2^ and absorbed the squared terms into the potential term *V* [*α*(*t*)] giving us an effective potential *V*_eff_ [*α*(*t*)]. We observe that Eq. (10) is the Euclidean action that models the effects of dissipation on a quantum-mechanical particle of mass *m*, with coordinates *α*(*t*), moving in a potential *V*_eff_ [*α*(*t*)], and subject to frictional forces with damping constant *η*′ ∝ *η* [17]. The slow prior Eq. (5) enforces only a local time constraint and maps to a kinetic energy term, whereas the environmental power spectrum prior Eq. (6) may give rise to a nonlocal time constraint and maps to a dissipative term.

The case of *n* = 1 is identical to the Caldeira-Leggett action [18] whereby the effect of ohmic dissipation on a quantum particle is modeled by linear coupling to an external heat bath consisting of an ensemble of harmonic oscillators. This functional-integral description of macroscopic quantum friction is of widespread interest in condensed matter physics and more recently in quantum computing where it provides a model of decoherence of a qubit [19]. In a metastable system, subohmic dissipation (0 < *n* < 1) can give rise to localization of a quantum particle, in contradistinction to the dispersed dynamics arising from dissipation (*n* > 2). Ohmic dissipation (n=1) exhibits criticality and the long-time behavior of a quantum particle depends crucially on the potential energy landscape *V*_eff_ and the phenomenological friction constant *η*. In terms of our Bayesian inference task this implies that the 1/f noise prior is special in that it can result in critical dynamics in the estimate of *α*(*t*).

To put flesh on a specific example we now explore the problem of optimally estimating *α* when there exist *C* = 2 possible interpretations which, for the sake of generality, may not be equally probable. An actual realization of this would be perception of the Necker cube where the contrast of the outline intentionally highlights one perceptual image over the other. This scenario is particularly interesting since the model estimation problem can be mapped to an Ising model with long-range interactions and an external field.

The potential energy term corresponding to bistable perception is a double-well potential with minima at 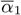 and 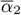. For convenience we take the curvatures at the two vacua 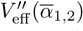 to be identical. A small bias in one percept is represented by a slight asymmetry in the potential (Figure 1). Naively we may expect that the optimal solution consists of *α*(*t*) localizing in one of the two wells, but since the mapping is to a quantum system, and not a classical system, we also have the possibility of delocalized solutions. In the undamped (*η* = 0) case we know that non-perturbative semiclassical solutions exist, representing instanton solutions whereby the estimate of *α* hops (quantum tunnels) from one well to another well in a short time 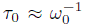 where 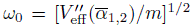 is the frequency of small oscillations in each well. In the limit that the barrier height is much greater than *ω*_0_ the trajectories of *α* can be described by an alternating sequence of hops from one well to another well. In the Bayesian inference problem these instantons are percept switches between two metastable percepts. Note that this implies that, with negligible environmental priors Eq. (6), the most likely (optimal) estimate of *α* is to always jump from one percept to another, and no neural noise is needed to elicit the percept switches.

**FIG. 1:**
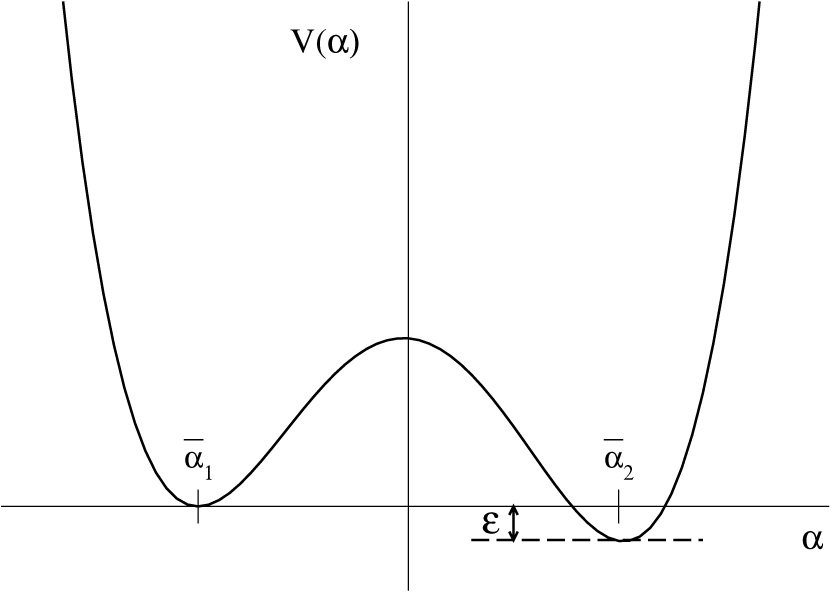
Potential energy landscape for *α* where there exist two valid interpretations of the data, one at 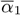 and the other 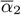. There is a small preference for the 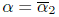 perception

By ignoring time-scales smaller than 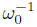 we can conveniently reduce the action to a two-state quantum system, where the eigenstates of the Pauli spin matrix *σ*_*z*_ corresponds to the two alternate percepts. The incorporation of dissipation, the second term in Eq. (10), into this approach leads to the well known spin-boson model, where many results are known [17]. In particular, in the case of ohmic dissipation (*n* = 1) there is a critical value of the dissipation constant 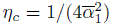, above which the particle is delocalized and below which the particle tunnels backwards and forwards.

A useful approximation is to discretize time in steps of *τ*_0_, and the dissipative contribution to the action then becomes

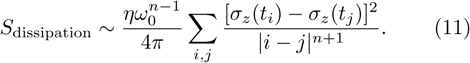

It is evident that we can identify the problem of estimating *α*(*t*) with a configuration of interacting Ising spins *s*_*i*_ where *i* indexes the time in units of *τ*_0_. Thus the dynamics of bistable perception with a power-law spectrum prior is equivalent to the instanton hops of a dissipative two-state system and is also equivalent to the spin configuration of a long-ranged one-dimensional Ising model (see Figure 2). The probability distribution for the spin sequence is then given by one-dimensional Ising model with power-law interactions

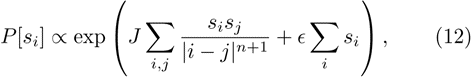

where 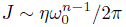. Using the correspondence between the dissipative two-state system and the Ising model with long range interactions, a number of properties rigorously proven for the Ising model can be directly transferred to the dynamics of bistable perception. Consider the unbiased case, (*ɛ* = 0). When *n* > 1, long-range order is absent in the Ising model and the percept always switches between two equally plausible percepts. For *n* < 1, symmetry breaking occurs and thus *α*(*t*) is localized. The case of *n* = 1 maps to the well-known inverse-square Ising model [20, 21], where there exists a phase transition demarcating the two types of behaviour at a critical value of *J*. Thus the non-local 1/f noise prior, reflecting the actual power spectra observed in the environment, gives rise to critical dynamics in the optimal estimate of *α*(*t*), and both stable perception and unstable perception are possible depending on the strength of the prior. To connect our results with experimental data we recall that perceptual alternations can be slowed and even stopped by intermittant removal of the visual stimulus [5], suggesting that memory (non-local) effects become important. Thus, when data is lacking, the brain places more strength on the 1/f prior, forcing a transition from unstable perception to stable perception.

**FIG. 2:**
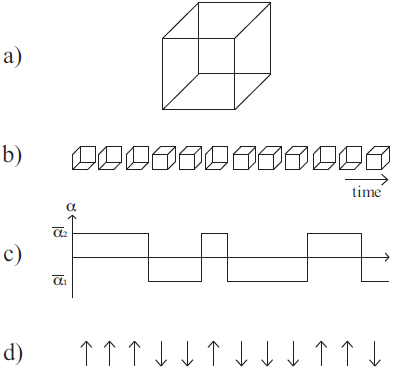
Equivalence between bistable perception switching, double-well quantum dynamics, and the Ising model. (a) Necker cube. (b) Switching between alternate percepts of the Necker cube. (c) Corresponding instanton trajectories between two stable model parameters. (d) Corresponding spin configuration of a one-dimensional 1/*r*^*n*+1^ Ising model.

This qualitative picture of localized/non-localized transitions does not change when there are *C* > 2 metastable states. The Caldeira-Leggett model of ohmic quantum dissipation has been studied when the potential energy landscape is taken to be sinusoidal (*C* = ∞), where it is known there again exists a critical value of the dissipation parameter *η* which separates a localized phase from a nonlocalized phase [22].

What happens when we include the effect of nonnegligible fluctuations in the input data? At any point in time, these fluctuations will tend to bias one model over the other. In the context of the Ising model the input noise, which we take to be white, can be modeled as quenched disorder. The Hamiltonian *H* for such a system is given by

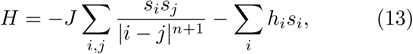

up to a temperature factor. The random field is taken to be normally distributed with a variance *σ* about the bias energy, i.e. 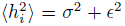. To determine whether symmetry breaking occurs in this random field Ising model we extend the elegant Imry-Ma argument [23] to the case of long-ranged interactions between the spins. Imagine we have an ordered Ising spin chain, aligned with the external field *ε*, and we then flip *L* of them where *L* ≫ 1. The energy cost from the change in spin-spin interactions will scale slower than log *L* for *n* > 1 and as *L*^1−*n*^ for 0 < *n* < 1. From the Central Limit Theorem the typical change in energy due to the external field will be 2*ɛL* − 2*σL*^1/2^. Whether the symmetry broken state is energetically favorable will thus depend on the relative values of *η*, *ε* and *σ*. In the unbiased system, the ordered state will be unstable for *n* > 1/2 and stable for *n* < 1/2 at low enough temperatures. When translated back to the bistable estimation problem these results predict that, when incorporating 1/*f* noise priors, a fluctuating bias in the input will prevent stable perception.

To summarize, the framework presented here shows that the combination of stimulus cues and internal representations of the physical world gives rise to the qualitative dynamics reported in multistable perception phenomena. This lends further weight to the compelling idea that the brain attempts to represent the entire posterior distribution rather than just a point estimate [24]. By mapping the statistical estimation task to a statistical physics problem we have shown how both stochastic switching and certainty may arise in the face of ambiguous signals. The non-local 1/f noise prior leads to critical dynamics of perception and allows a specific model to be learnt amongst a plethora of possible models.

Multistable perception has also been been reported in other sensory modalities, such as audition, olfaction and touch [4], suggesting that it may arise from a general neural design principle. The principles of probabilistic Bayesian inference and statistical mechanics provide a way to address perception dynamics whilst evoking very few assumptions about the sensory system.

Finally, the work presented here strengthens the connection between estimation theory and statistical mechanics, permitting us to understand the collective emergence of stochasticity and certainty more generally in statistical learning theory as well as perception dynamics.

J. Kinney and A. Churchland are acknowledged for helpful discussions and comments on the manuscript. This work was supported by the Simons Foundation.

## References

[1] D. M. Eagleman, Nature 2, 920 (2001).

[2] R. Blake and N. Logothetis, Nat. Rev. Neuro. 3, 1 (2002).

[3] L. A. Necker, London and Edinburgh Philosophical Magazine and Journal of Science 3, 329 (1832).

[4] J. Schwartz, N. Grimault, J. Hupe, B. C. J. Moore, and D. Presnitzer, Phil. Trans. R. Soc 367, 896 (2012).

[5] D. A. Leopold, M. Wilke, A. Maier, and N. K. Logothetis, Nature Neuroscience 5, 605 (2002).

[6] A. P. Murphy, D. A. Leopold, and A. E. Welchman, Frontiers in Psychology 5, 60 (2014).

[7] M. Riani and E. Simonotto, Phys. Rev. Lett. 72, 3120 (1994).

[8] R. Moreno-Bote, J. Rinzel, and N. Rubin, Journal of Neurophysiology 8(7), 1 (2007).

[9] G. Huguet, J. Rinzel, and J. Hupe, Journal of Vision 14(3):19, 1 (2014).

[10] W. Bialek and M. DeWeese, Phys. Rev. Lett. 74, 3079 (1995).

[11] G. S. Atwal and W. Bialek, in Advances in Neural Information Processing Systems, edited by S. Thrun, L. Saul, and B. Scholkopf (MIT Press, Cambridge CA, 2004), vol. 16.

[12] T. M. Cover and J. A. Thomas, Elements of Information Theory (Wiley-Interscience, 2006), 2nd ed.

[13] R. F. Voss and J. Clarke, Nature 258, 317 (1975).

[14] M. B. Weissman, Rev. Mod. Phys. 60, 537 (1988).

[15] P. C. Ivanov, M. G. Rosenblum, Z. R. S. L. A. Nunes Amaral, S. Havlin, A. L. Goldberger, and H. E. Stanley, Nature 399, 461 (1999).

[16] P. Bak, C. Tang, and K. Wiesenfeld, Phys. Rev. Lett. 59, 381 (1987).

[17] A. J. Leggett, S. Chakravarty, A. T. Dorsey, M. P. A. Fisher, A. Garg, and W. Zwerger, Rev. Mod. Phys. 59, 1 (1987).

[18] A. O. Caldeira and A. J. Leggett, Phys. Rev. Lett. 46, 211 (1981).

[19] M. A. Nielsen and I. L. Chuang, Quantum Computation and Quantum Information (Cambridge University Press, 2011).

[20] D. J. Thouless, Phys. Rev. 187, 732 (1969).

[21] P. W. Anderson and G. Yuval, J. Phys. C 4, 607 (1971).

[22] M. P. A. Fisher and W. Zwerger, Phys. Rev. B 32, 6190 (1985).

[23] Y. Imry and S. Ma, Phys. Rev. Lett. 35, 1399 (1975).

[24] S. J. Gershman, E. Vul, and J. B. Tenenbaum, Neural Computation 24, 1 (2012).

